# The Rcs stress response system modulates *Serratia marcescens* induced inflammation and bacterial proliferation in a rabbit keratitis model

**DOI:** 10.1101/2020.08.07.242446

**Authors:** Eric G. Romanowski, Nicholas A. Stella, John Romanowski, Kathleen A. Yates, Deepinder K. Dhaliwal, Robert M. Q. Shanks

## Abstract

In this study, we tested the hypothesis that the conserved bacterial Rcs stress response system mediates corneal pathogenesis associated with *Serratia marcescens* ocular infections. This was accomplished by modifying Rcs activity using mutant strains. These include a mutant that has a hyper-active Rcs system due to deletion of the IgaA family gene, *gumB*, and a *gumB rcsC* double mutant that is defective for Rcs signaling. The role of the Rcs system and bacterial stress response systems for microbial keratitis is not known. Here we observed that the Rcs-activated *gumB* mutant had a >50-fold reduction in proliferation compared to the wild type within rabbit corneas at 48 h, and demonstrated a notable reduction in inflammation based on inflammatory signs and proinflammatory markers measured at the RNA and protein levels. The *gumB* mutant phenotypes could be complemented by wild-type *gumB* on a plasmid and partially complemented by restoration of *shlA* cytolysin expression and elimination of capsular polysaccharide production. We observed that inactivation of the Rcs stress response system completely restored corneal virulence to the *gumB* mutant. NanoString transcriptional analysis of bacterial genes expressed during microbial keratitis demonstrated expression of *gumB, rcsB, shlA*, and three metalloprotease genes. Data suggest that the bacterial capsular polysaccharide is not necessary for infection, but capsule overexpression reduces inflammation. Together, these data indicate that GumB regulates virulence factor production through the Rcs system and this overall stress response system is a key mediator of a bacterium’s ability to induce vision-threatening keratitis.

## INTRODUCTION

The role for bacterial stress systems has been demonstrated as being essential for establishing wild-type levels of microbial pathogenesis in several animal models of infection (1). These include bacterial cell envelope stress response systems in both Gram-negative and Gram-positive bacteria that protect the cell envelope from external and internal host-defense factors (2). The Rcs system is well-studied in *Eschericia coli* and *Salmonella enterica* and conserved in other members of the Enterobacteriaceae (3). In *E. coli* and *S. enterica*, the Rcs system has a role in responding to envelope stress caused by membrane affecting agents including detergents and cell wall damaging compounds such as lysozyme (3). Mutational inactivation of the Rcs system has been reported to increase the virulence potential of several bacteria including *Proteus mirabilis* in a *Galleria mellonella* infection model (4), *Edwardiella tarda* in a zebrafish model (5), and *S. marcescens* in a murine bacteremia model (6). Similarly, the increased virulence of *Yersinia pestis* compared to other *Yersinia* species is thought to be partially due to an incomplete Rcs system (3). In other models, Rcs system-defective mutants are less virulent. For example, *Citrobacter rodentium* and *E. coli rcsB* mutants were less virulent in rodent intestinal infection models (7, 8). Therefore, it is likely that bacteria tightly regulate the Rcs system during infection because it is important in regulating the expression of genes that balance host-pathogen interactions.

The Rcs system is inhibited by the inner-membrane protein, IgaA, and is essential for growth in *E. coli* and *Salmonella* species (3). The *S. marcescens* IgaA ortholog, GumB, is functionally conserved with other IgaA-family proteins, as *gumB* mutant defects can be complemented in trans by the *E. coli, Klebsiella pneumoniae*, and *S. enterica* IgaA orthologs (9). However, unlike *E. coli* and *S. enterica*, IgaA-proteins are not essential for viability in *P. mirabilis* (10) or *S. marcescens* (9). It is also likely not essential in *Y. pestis*, as it has not turned up in genome wide screens for essential genes (3). Evidence from previous studies suggests that loss of *gumB* locks *S. marcescens* in a “stressed-out” state leading to phenotypes consistent with an activated stress response including loss of flagella production, reduced secretion of toxins, and excess capsular polysaccharide synthesis (9, 11). These phenotypes are reminiscent with those of bacteria in a chronic infection or in biofilms and are associated with a less acute infection (12, 13). Consistently, mice infected with *S. enterica* that coded for partial function IgaA protein were severely defective in systemic mouse infection models (14).

Although mutation of genes for IgaA-family proteins generally reduces virulence, there is reason to suggest that in some circumstances inactivation of IgaA may increase bacterial persistence and resistance to the immune system. For example, in an *in vitro* experiment in which *E. coli* was serially passaged with phagocytic cells, the surviving *E. coli* gained mutations in the *igaA* gene that increased capsule production and resistance to phagocytosis (15). Mutations in *igaA* have also been characterized among multidrug resistant clinical isolates of *E. coli* (16), together suggesting that mutation of *igaA* can aid in pathoadaptation. Consistently, the *igaA* gene was originally found in a screen for mutations that enabled *S. enterica* serovar Typhimurium to proliferate within fibroblast cells (17).

Because of the importance of IgaA-family protein GumB in *S. marcescens* proliferation in an invertebrate infection model, regulation of virulence factors, and cytotoxicity to corneal cells *in vitro* (9, 11), we tested the prediction that GumB and the Rcs system is crucial for corneal infections. In this study, we analyze the importance of the Rcs system in *S. marcescens* keratitis using a rabbit keratitis model and measure gene expression of known and suspected virulence factor genes *in vivo*. Together, this study indicates that over activation of a bacterial stress response system and the associated shift to a chronic rather than acute-infection lifestyle remarkably reduces pathogenesis without strongly affecting bacterial proliferation.

## RESULTS

### GumB is necessary for *S. marcescens* proliferation and inflammation in a rabbit **corneal** infection model

*S. marcescens* is a leading cause of contact lens associated corneal infections (18, 19). Previous results using a keratitis isolate of *S. marcescens* indicated that constitutive activation of the Rcs stress response system, through mutation of the *gumB* gene, dramatically reduces the bacterium’s ability to cause cytotoxicity *in vitro* using primary human corneal cells and virulence in an invertebrate pathogenesis model (11). However, the influence of the Rcs system on bacterial pathogenesis in a keratitis model is unknown. Cultures of *S. marcescens* wild-type contact lens-associated keratitis isolate K904 (WT) and an isogenic Δ*gumB* mutant were grown overnight in LB medium, diluted in PBS, and ∼500 CFU in 25 µl were injected into the corneal stroma. The actual injection was 468±297 for WT and 885±550 for Δ*gumB*.

At 24 and 48 hours post-inoculation, corneal and conjunctival clinical signs were examined by slit-lamp and evaluated using parameters from a modified MacDonald-Shadduck scoring system (20). At 24 hours post-inoculation, there was little difference in clinical inflammation scores for the cornea (Figure 1A,B); however, the infiltrates were distinct. Corneas infected with the WT demonstrated a snow-flake like infiltrate in 11 out of 12 eyes, whereas the Δ*gumB* mutant infected corneas had globular infiltrate in 11 out of 11 corneas (Figure 1A,C).

**Figure 1.**
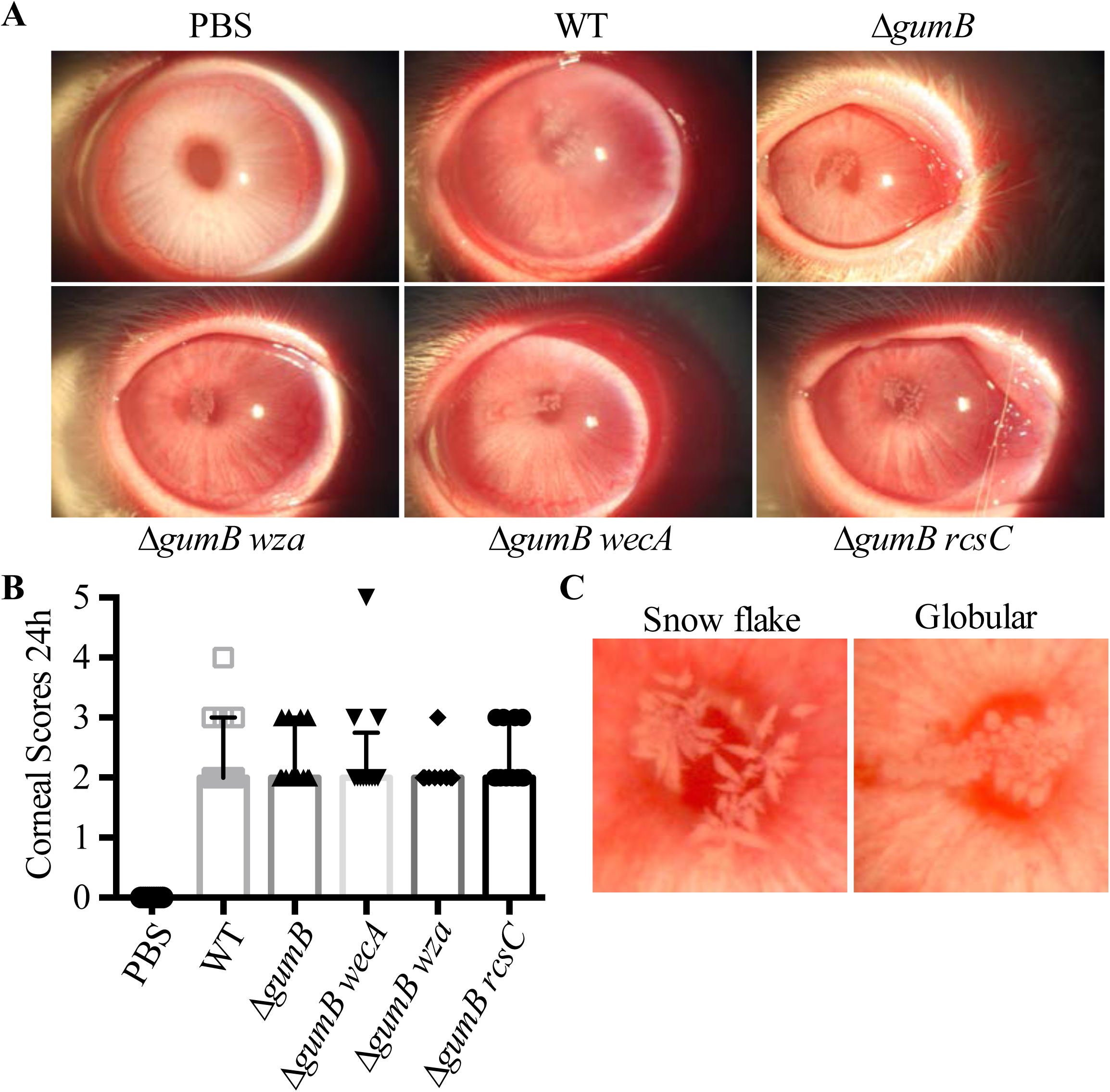
Rcs status of bacteria influences infiltrate morphology during *S. marcescens* corneal infections. **A**. Representative images of infected eyes at 24 h post-infection. **B**. Median values and interquartile ranges for clinical signs of inflammation 24 h after infection. **C**. Representative images of typical infiltrate types.

At 48 hours, corneal inflammation was higher for eyes infected with the WT than the Δ*gumB* mutant (Figure 2). The corresponding corneal inflammatory scores were significantly higher (p<0.001, Kruskal-Wallis with Dunn’s posttest) for the WT (median 8.0, 95% CI = 7.13-8.21) compared to the Δ*gumB* mutant (median 3.0, 95% CI = 2.44-4.36) (Figure 3A). Corneal scores were the culmination of 0-4 for corneal opacity, area of corneal opacity, corneal vascularization, and corneal staining with fluorescein. Injection of PBS alone as a negative control resulted in a score of 0±0 (Figure 2, 3A). Rabbit eyes infected with WT *S. marcescens* presented with medium to large central corneal ulcers with fluorescein staining indicating loss of the epithelium (Figure 2 and 3B). The Δ*gumB* mutant infected eyes either had small ulcers or no ulcer. However, the epithelial defect area did not reach significance between the WT and Δ*gumB* mutant.

**Figure 2.**
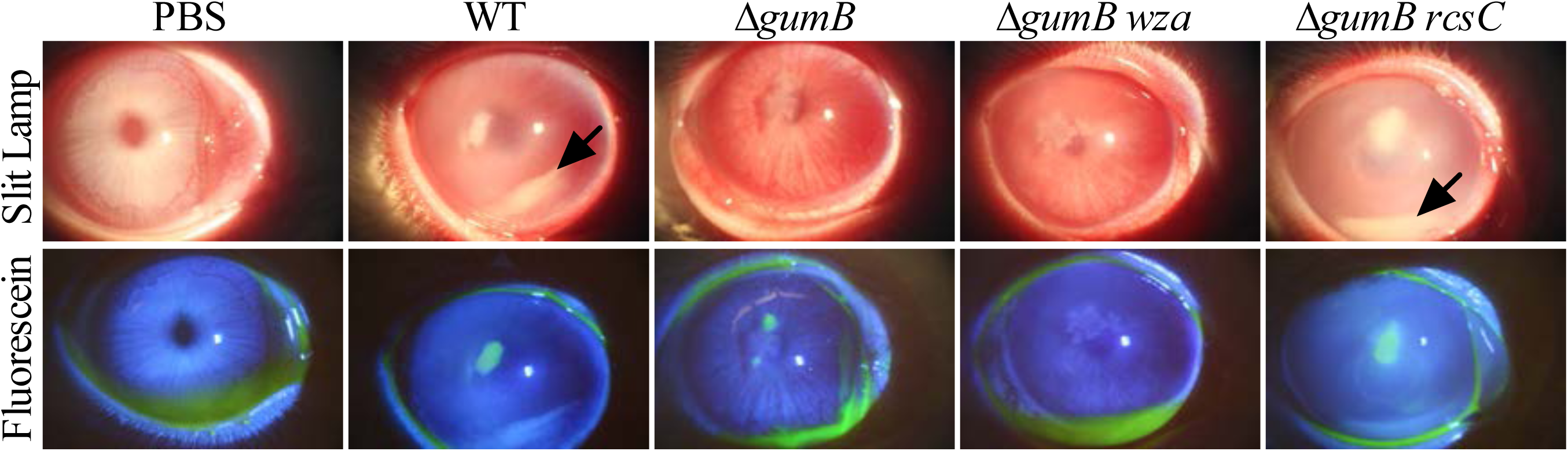
GumB is required for wild-type inflammation during *S. marcescens* corneal infections. Representative images of infected eyes at 48 h post-infection. Fluorescein staining represents epithelial damage. The arrows indicate hypopyons.

**Figure 3.**
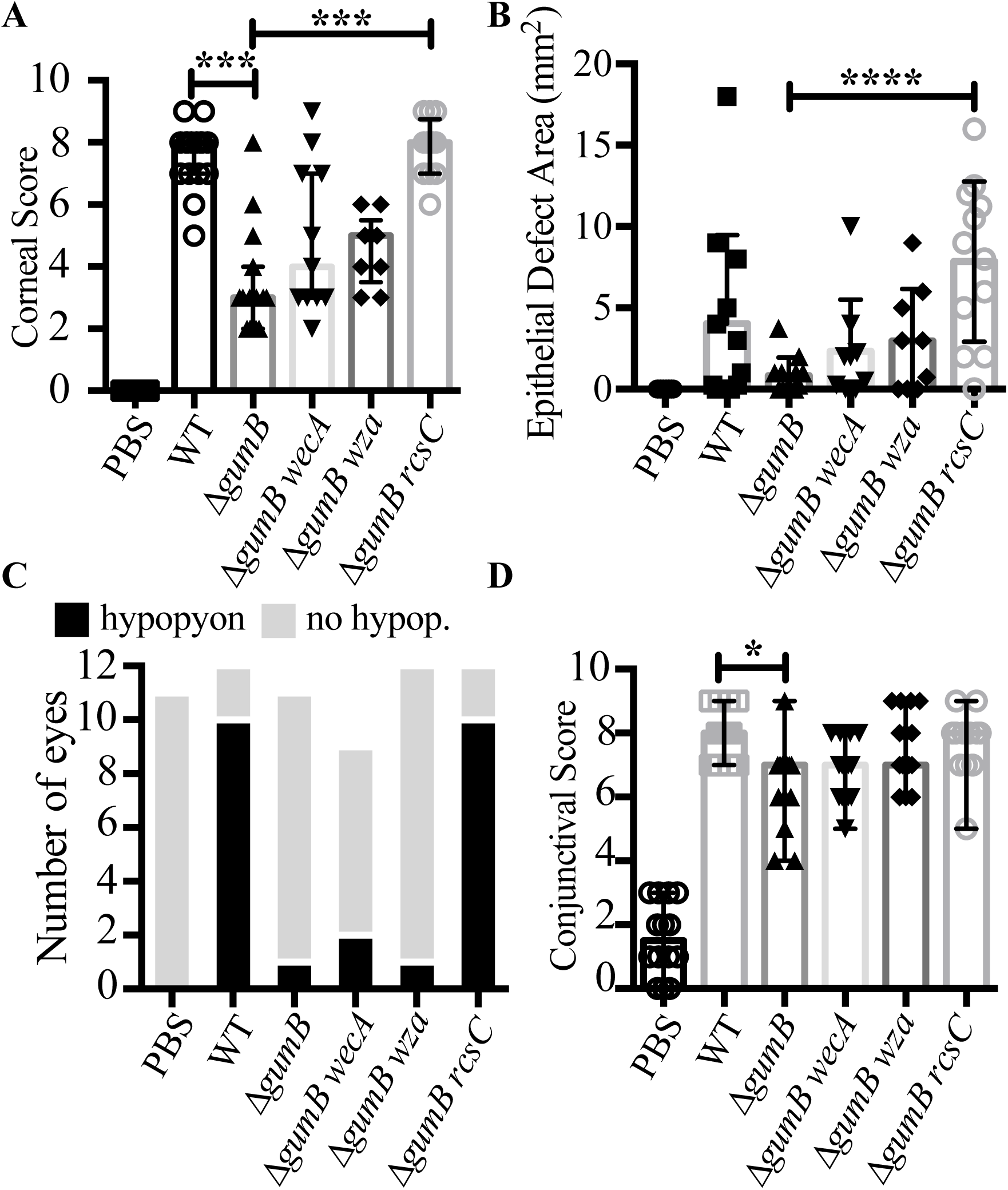
GumB and the Rcs system dictate the ability of *S. marcescens* to induce inflammation during corneal infection. Median values and interquartile ranges for clinical signs of inflammation 48 h after infection for **A**. cornea and **D**. conjunctiva. The asterisks indicate significances by Kruskal-Wallis with Dunn’s post-test. *=p<0.05, ***=p<0.001, ****=p<0.0001. **B**. Average and standard deviation of epithelial defects. ****= p<0.0001, ANOVA with Tukey’s post-test. **C**. Enumeration of eyes with hypopyon.

Other inflammatory signs included as loss of corneal clarity and the presence of hypopyons, a build-up of inflammatory cells in the anterior chamber (Figure 2). Hypopyons are indicative of a high level of ocular inflammation and predictive of worse clinical outcomes (21, 22). Notably, 9 out of 12 eyes infected with the WT demonstrated a hypopyon (Figure 2 arrow) compared to 0 out of 17 for PBS injected eyes (p=0.0002, Fisher’s Exact test) and 1 out of 11 for Δ*gumB* infected eyes (p=0.0028 compared to WT) (Figure 3C and Table 2). Conversely, eyes infected with the Δ*gumB* mutant were generally clear except for globular infiltrates (Figure 1A, 2).

**Table 1.**
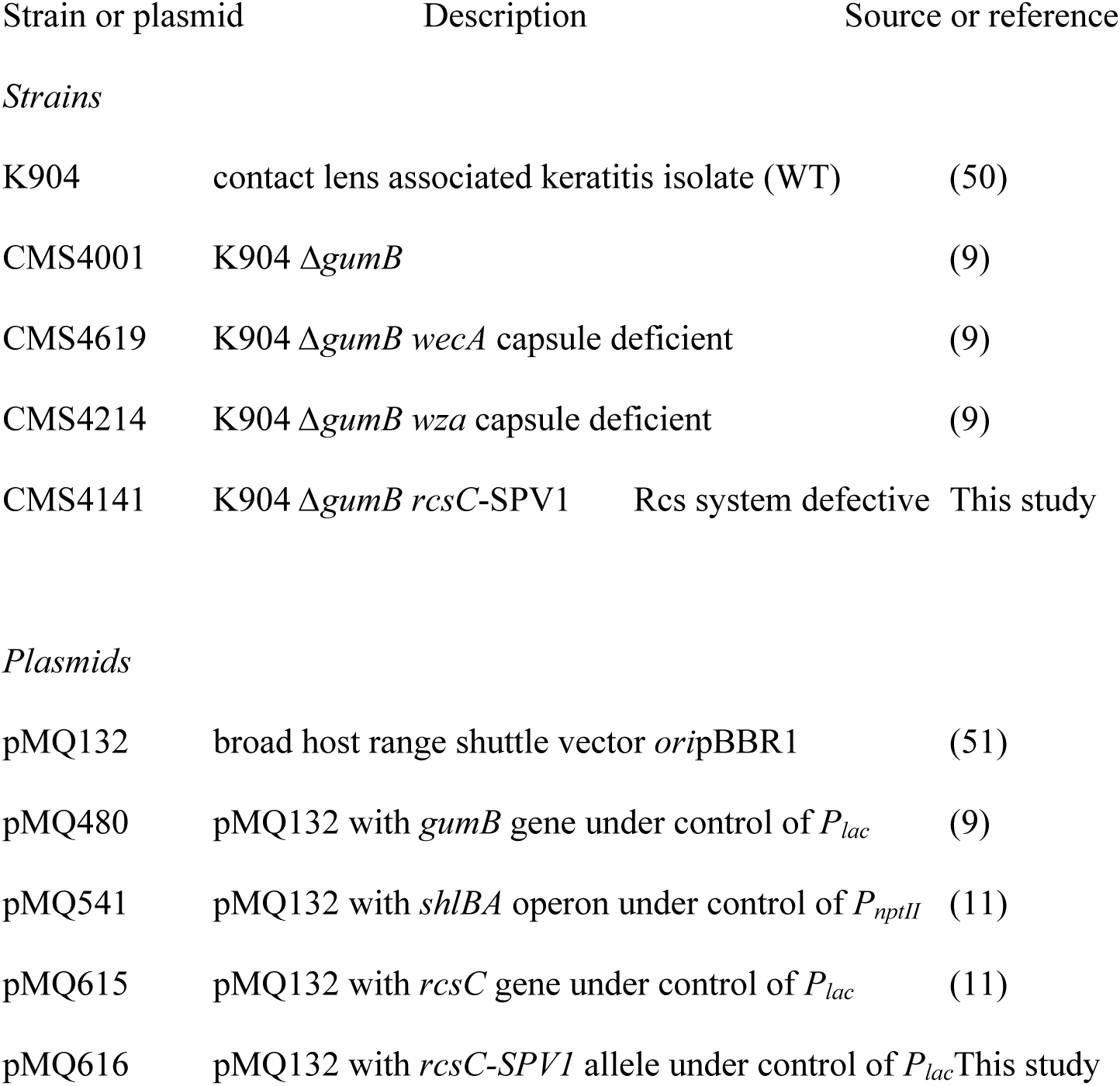
Strains and plasmids used in this study.

| Strain or plasmid | Description | Source or reference |
| --- | --- | --- |
| *Strains* |  |  |
| K904 | contact lens associated keratitis isolate (WT) | (50) |
| CMS4001 | K904 $\Delta gumB$ | (9) |
| CMS4619 | K904 $\Delta gumB$ *wecA* capsule deficient | (9) |
| CMS4214 | K904 $\Delta gumB$ *wza* capsule deficient | (9) |
| CMS4141 | K904 $\Delta gumB$ *rscC*-SPV1 Rcs system defective | This study |
| *Plasmids* |  |  |
| pMQ132 | broad host range shuttle vector *ori*pBBR1 | (51) |
| pMQ480 | pMQ132 with *gumB* gene under control of $P_{lac}$ | (9) |
| pMQ541 | pMQ132 with *shlBA* operon under control of $P_{nptII}$ | (11) |
| pMQ615 | pMQ132 with *rscC* gene under control of $P_{lac}$ | (11) |
| pMQ616 | pMQ132 with *rscC*-SPV1 allele under control of $P_{lac}$ | This study |

**Table 2.** Hypopyons were less prevalent in Δ*gumB* infected eyes.

|  | **Hypopyon<sup>a</sup>** | **No**<br>**Hypopyon<sup>b</sup>** | **P-value vs WT<sup>c</sup>** | **P-value vs PBS<sup>c</sup>** |
| --- | --- | --- | --- | --- |
| PBS | 0 | 11 | **0.0001** | NA |
| WT | 10 | 2 | NA | **0.0001** |
| $\Delta gumB$ | 1 | 10 | **0.0006** | 1.0000 |
| $\Delta gumB\ wecA$ | 2 | 7 | **0.0092** | 0.1895 |
| $\Delta gumB\ wza$ | 1 | 11 | **0.0006** | 1.0000 |
| $\Delta gumB\ rcsC$ | 10 | 2 | 1.0000 | **0.0001** |

<sup>a-b</sup> Number of eyes with or without hypopyons or extensive fibrin in the anterior chamber.
<sup>c</sup> Two tailed Fisher's Exact test, NA = not applicable, significant differences in bold text.

Conjunctival inflammatory scores followed a similar pattern to corneal scores with WT infected eyes having significantly higher scores than the *ΔgumB* bacteria (p<0.05, Kruskal-Wallis with Dunn’s posttest), although the magnitude of the difference between WT and Δ*gumB* infected eyes was less extensive than for corneal inflammation. Scores for both were higher than PBS-injected eyes (p<0.0001 for WT and p<0.05 for Δ*gumB*) (Figure 3D).

We assessed pro-inflammatory cytokines IL-1β and Tnfα at the RNA level by NanoString technology in infected corneas at 24 hours post-inoculation (Fig 4A). Relative to the PBS injected eyes, and normalized by GAPDH levels, Tnfα transcript were up 4.9 fold in WT infected eyes compared to PBS infected eyes and 5.7 fold compared to Δ*gumB* mutant infected eyes (p<0.05 by Mann-Whitney). The effect was more dramatic for IL-1β with a 363.8-fold increase in WT injected eyes compared to PBS injected, and 26.2-fold higher in the WT infected eyes than Δ*gumB* infected eyes (p<0.05, Mann-Whitney) (Figure 4A). Because the effect was greater for IL-1β, we evaluated IL-1β protein levels 48h post-inoculation by ELISA. A >1000-fold increase in IL-1β was measured in WT infected eyes compared to PBS infected eyes (p<0.01, ANOVA with Tukey’s posttest) (Figure 4B). Eyes infected with Δ*gumB* had a 56-fold elevation in IL-1β compared to PBS, but this did not reach significance. The Δ*gumB* mutant corneas had 19-fold reduced IL-1β compared to WT infected corneas (p<0.01) (Figure 4B).

**Figure 4.**
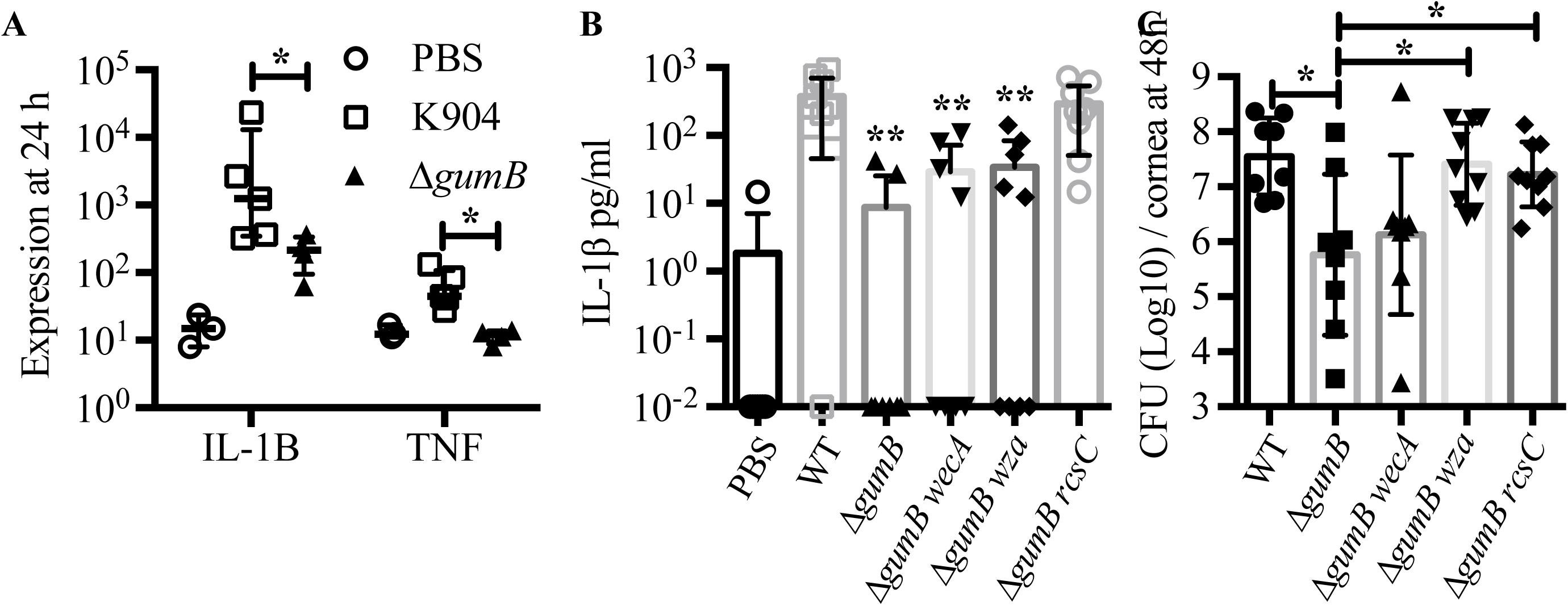
GumB mediates inflammation and proliferation within the rabbit cornea. **A**. Median values and interquartile ranges are shown of RNA transcripts normalized by GAPDH RNA levels from corneas 24 hours after infection. The asterisk indicates p<0.05 WT versus Δ*gumB* by Mann-Whitney test (n=3-5 per group). **B**. Mean and standard deviation of IL-1B protein levels measured by ELISA 48 h post infection (n=8-9 per group). ** = p<0.01 compared to WT using ANOVA with Tukey’s post-test. **C**. Mean and standard deviation of CFU per cornea at 48 h post infection. Asterisks indicate p<0.05 differences by ANOVA with Tukey’s test.

The bacterial burden, at 48 hours post-infection, was 61-fold higher in the wild type infected corneas compared to those infected by the Δ*gumB* mutant (Figure 4C), p<0.05, ANOVA with Tukey’s posttest. Consistent with higher bacterial numbers and inflammatory scores, hematoxylin and eosin (H&E) staining of corneal sections showed epithelial defects, edema, and neutrophils in the wild type infected corneas (Figure 5). By comparison Δ*gumB* infected corneas had intact epithelial layers and mild edema. Notably, the Δ*gumB* mutant infected corneas had globular infiltrates with a biofilm-like morphology (Figure 5).

**Figure 5.**
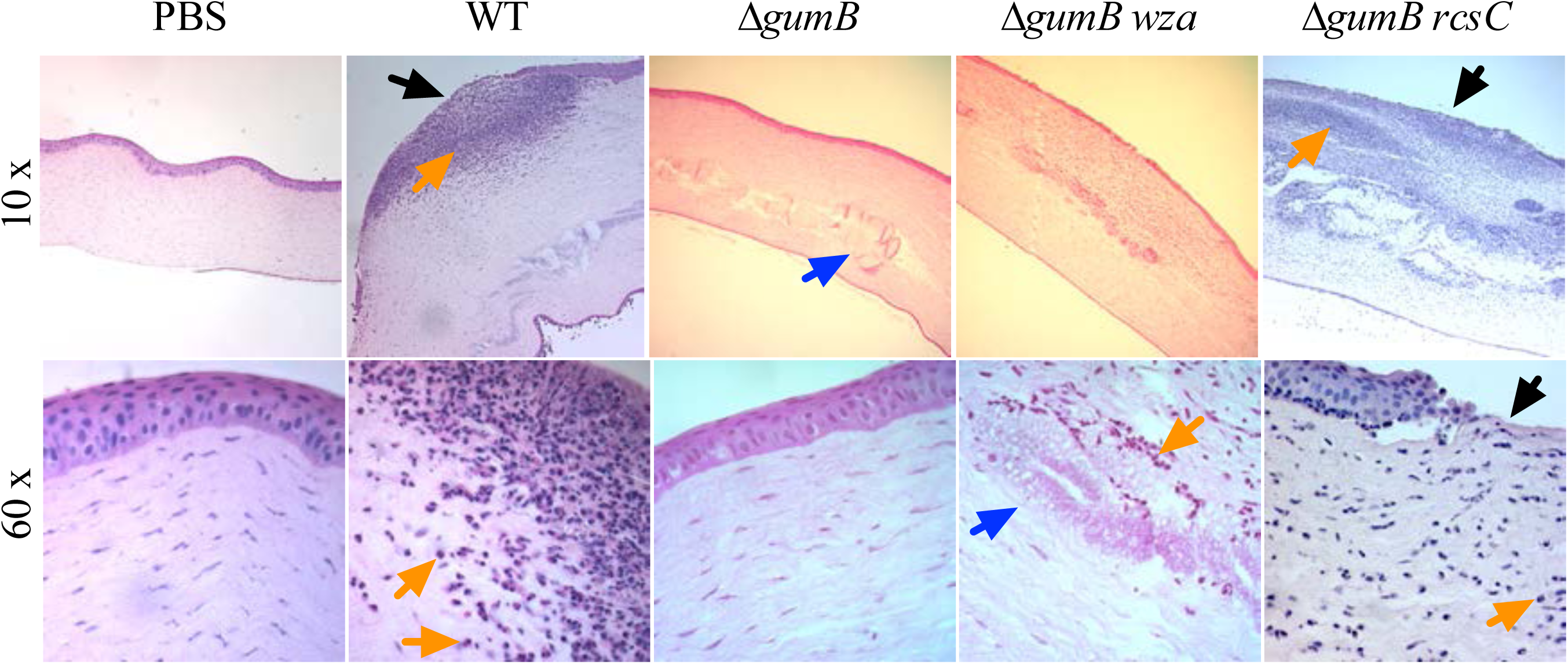
Histological analysis of *S. marcescens* infected corneas at 48 h. Representative images of hemotoxylin and eosin stained corneal sections of eyes at 48 h post-infection. Large neutrophil infiltrates were observed in the WT and Δ*gumB rcsC* infected eyes. The objective magnification is indicated. Black arrows indicate epithelial defects; blue arrows indicate globular infiltrates in Δ*gumB* mutant-infected corneas; orange arrows indicate neutrophils.

### The Δ*gumB* mutant inflammatory defect can be complemented by wild-type *gumB* on a plasmid

Complementation analysis was performed to ensure that the Δ*gumB* defect was due to mutation of *gumB* and not another unknown defect in the Δ*gumB* mutant strain or polar effect of the deletion (Figure 6). Similar to the analysis above, the WT with an empty vector negative control injected with an average of 349±11 CFU per cornea induced median corneal inflammatory scores of 7.0 (n=8, 95% CI = 4.89-8.11) at 48 h post-inoculation (Figure 6A). The Δ*gumB* mutant with the empty vector negative control injected with 558±90 CFU per cornea caused inflammatory scores of 2.0 (n=8, CI = 1.71-5.04). Eyes were more inflamed, with a score of 7.0 (n=7, CI = 5.70-7.16) when infected with the Δ*gumB* mutant with wild-type *gumB* on a plasmid (370±36 CFU injected per cornea) (Figure 6A), and were similar to the WT injected corneas. PBS injected corneas induced corneal scores of 0 (n=8, 95% CI = −0.34-0.84), and the WT + vector and Δ*gumB* + p*gumB* (wild-type *gumB* on a plasmid), but not Δ*gumB* with vector alone were significantly different than the PBS injection group (p<0.001, Kruskal-Wallis with Dunn’s posttest). A similar pattern was observed with conjuntival inflammatory scores where, all groups except the Δ*gumB* with empty vector group had significantly higher conjunctival inflammatory scores compared to PBS injected eyes (Figure 6B). Images of infected eyes confirm the relatively decreased inflammation and increased clarity of corneas infected with the Δ*gumB* mutant compared to the other groups (Figure 6C).

**Figure 6.**
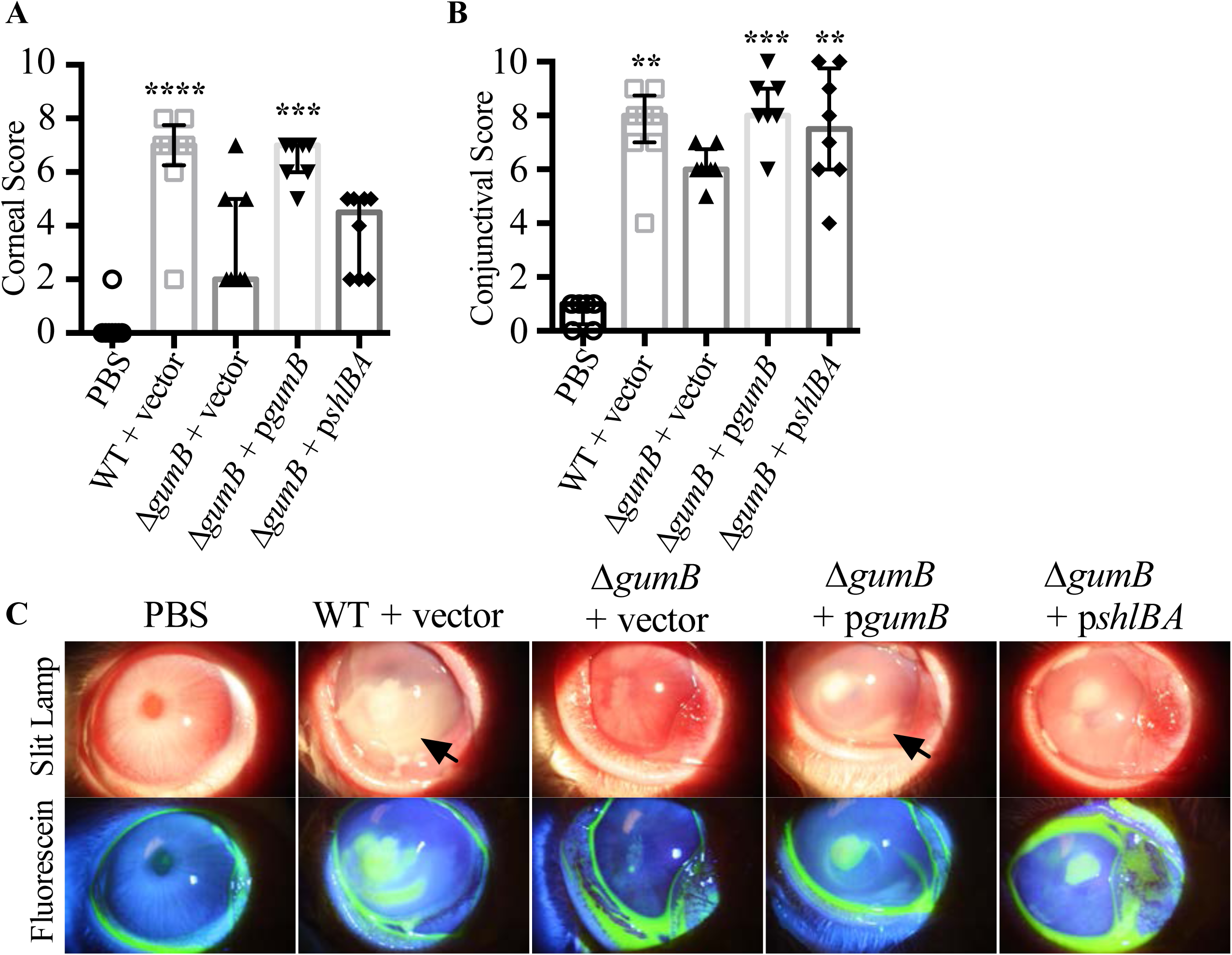
Complementation of Δ*gumB* and multicopy expression of cytolysin gene *shlA*. Median values and interquartile ranges are shown for corneal (A) and conjuntival (B) inflammatory scores at 48 hours after infection. Significant differences from the ΔgumB + vector group are indicated by asterisks as determined by Kruskal-Wallis analysis with Dunn’s post-test. *=p<0.05, ***=p<0.001, ****=p<0.0001, n=7-8 per group. **C**. Representative images of infected eyes at 48 h post-infection. Fluorescein staining represents epithelial damage. Black arrows indicate hypopyons.

The wild-type strain with the empty vector proliferated to 2.2 × 10^7^ CFU per cornea in 48 h compared to 2.7 ×10^6^ CFU for the Δ*gumB* mutant with the empty vector, a 8-fold reduction that did not reach significance (n=8), (p>0.05 by Kruskal-Wallis with Dunn’s multiple comparison test). With the *gumB* plasmid in the Δ*gumB* mutant CFU per cornea increased to 1.4 ×10^7^ (n=7), which was indistinguishable from the wild type (Figure 7A).

**Figure 7.**
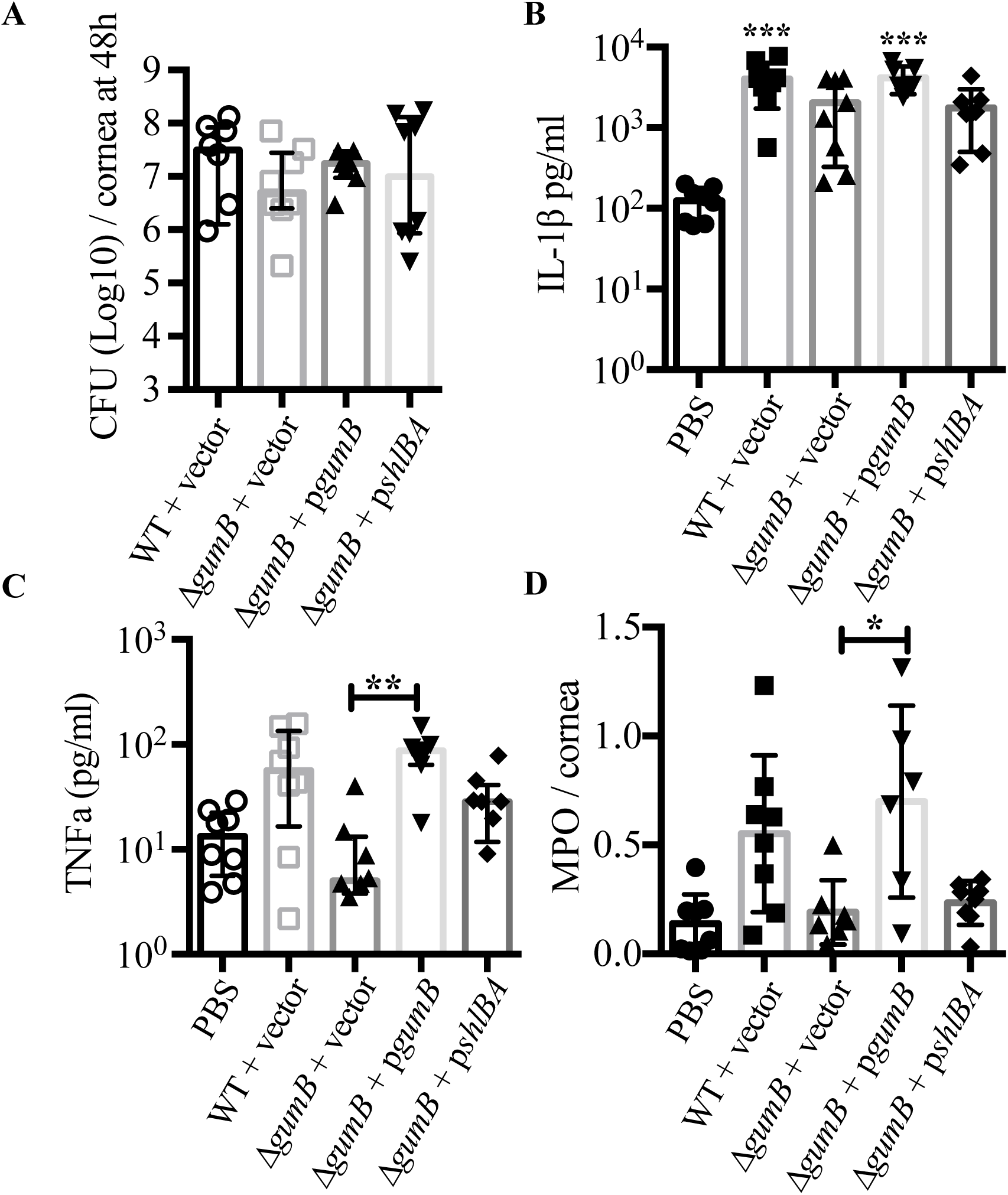
GumB mediates inflammation proliferation within the rabbit cornea. **A**. Mean and standard deviation CFU values per cornea for 48 hours after infection. **B-C**. Mean and standard deviation of IL-1B (**B**) and TNFa (**C**) protein levels measured by ELISA 48 h post infection (n=7-8 per group). *** = p<0.001 compared to the PBS group by ANOVA with Tukey’s post-test for B, and **=p<0.01. **D**. Mean and standard deviation of MPO levels per cornea at 48 h post infection. Asterisks indicate p<0.05 differences by ANOVA with Tukey’s test.

Corneal IL-1β levels were reduced approximately 2-fold from 4032±2308 pg/ml in WT with the empty vector control compared to the Δ*gumB* mutant with the vector control (2051±1725 pg/ml). PBS injected eyes had 124±55 pg/ml. The Δ*gumB* mutant with *gumB* on a plasmid had 4188±1582 pg/ml. (Figure 7B). A similar trend was observed with Tnfα (Figure 7C).

Since neutrophil mediated damage is a major cause of corneal pathology during microbial keratitis we indirectly measured neutrophil infiltration by biochemically measuring myeloperoxidase (MPO) activity. A clear increase in MPO activity was measured from corneas infected with the wild-type bacteria with vector compared to PBS injected corneas (Figure 7D), p<0.05, by ANOVA with Tukey’s posttest. The Δ*gumB* mutant with the same vector was defective in attracting neutrophils and was not different from the PBS-injected control group, p>0.05. Conversely, when the wild-type *gumB* gene was added back to the Δ*gumB* mutant, MPO activity was restored and significantly higher than the PBS group (p<0.01).

### Role of capsule and ShlA cytolysin in GumB mediated corneal infections

Mutation of IgaA-family proteins like *gumB* is noted to cause a major increase in production of extracellular polysaccharide (9). Here we used previously described isogenic variants of the *gumB* mutant that do not make capsule due to mutations in capsule biosynthetic genes *wecA* and *wza* (9) to test for the importance of over production of capsule in the reduced corneal inflammation phenotype of the Δ*gumB* mutant. At 24 hours the *gumB wecA* double mutant produced inflammation indistinguishable from the other groups (Figure 1), but at 48 h Δ*gumB wza* and Δ*gumB wecA* mutants induced intermediate clinical corneal scores that were elevated as compared to the Δ*gumB* mutant and significantly higher corneal inflammatory scores than PBS injected eyes (p<0.05), whereas the Δ*gumB* mutant infected eyes were not significantly different from PBS injected eyes (p>0.05, by Kruskal-Wallis with Dunn’s posttest) (Figure 3A). Epithelial defects and the frequency of hypopyons were similar to the Δ*gumB* mutant (Figure 3B-C). IL-1β levels for the *gumB wecA* mutant were intermediate compared to the Δ*gumB* and wild-type (Figure 4B). CFU were significantly elevated in the Δ*gumB wza* double mutant compared to the Δ*gumB* mutant (Figure 4C). H&E staining of Δ*gumB wza* double mutant eyes were similar to Δ*gumB* infected eyes, but showed intermediate levels of neutrophils and a less robust epithelial layer (Figure 5). Together, these suggest that over production of the capsule plays a role in the Δ*gumB* mutant keratitis defects, but it is not the only factor involved.

A previous study demonstrated that the Δ*gumB* mutant fails to transcribe and produce the ShlA cytolysin (11). The *shlA* cytolysin gene is expressed in an operon with the ShlA transporter gene *shlB*. To test whether lack of ShlA biosynthesis was the reason for the Δ*gumB* mutant defect, we expressed the *shlBA* operon from a plasmid in the Δ*gumB* mutant that was previously shown to complement the Δ*gumB* mutant cytotoxicity defective phenotype (11). The corneal inflammatory score went from 2.0 for the Δ*gumB* mutant with the vector to 4.5 for the Δ*gumB* mutant with the *shlBA* plasmid, compared to 7 for the wild type with vector (n=8) (Figure 6A, C). A similar pattern was observed for conjunctival signs of inflammation (Figure 6B, C). However neither of these changes reached significance. Expression of *shlBA* also partially restored bacterial proliferation in the cornea, from 5 ×10^6^ for the *gumB* mutant with the vector to 1 × 10^7^ for the *gumB* mutant with the *shlBA* plasmid (compared to 3 × 10^7^ for the wild type with the vector, p>0.05, ANOVA with Tukey’s posttest), (Figure 7A). IL-1β and MPO levels were not clearly modified by the *shlBA* plasmids (Figure 7A,C).

The relatively mild impact of the *shlBA* plasmid may come from plasmid instability. Whereas the same plasmid back-bone was used for the vector, *gumB*, and *shlBA* plasmids, there was a notable difference in recovery of bacteria with each of the plasmids after 48 hours in the rabbit cornea. After 48 h, 0% of the bacteria had the *shlBA* plasmid, compared to 76% for the vector and 81% for the *gumB* plasmid. Despite loss of the *shlBA* plasmid in the rabbit by 48 h, ectopic expression of *shlBA* conferred intermediate phenotypes that suggest at a least minor role for ShlA in corneal infections.

### The Δ*gumB* mutant keratitis phenotype is reversed through inactivation of the Rcs pathway

Data indicates that the Δ*gumB* mutant has an overactive Rcs system and that mutation of *rcsB* or *rcsC* genes restore wild-type like phenotypes to the Δ*gumB* mutant such as ShlA-dependent cytotoxicity to corneal cells (11). This suggests that over-activation of the Rcs system may be responsible for the reduced virulence of the Δ*gumB* mutant. In order to test whether the Rcs system is necessary for the Δ*gumB* mutant keratitis phenotype, we used a double mutant defective with a null mutation in the *rcsC* gene, a necessary component of Rcs signaling. We isolated a Δ*gumB* mutant with a spontaneous duplication of 49 bp in the *rcsC* gene that creates an out-of-frame mutation. This allele was named *rcsC*-SPV1, for <u>s</u>pontaneous <u>p</u>igmented <u>v</u>ariant number 1, and genetic experiments support that it codes for a null mutation (data not shown).

This Δ*gumB rcsC* double mutant was introduced into rabbit corneas (503±448 CFU average inocula) to test whether over-activation of the Rcs system due to *gumB* mutation compared to the WT was responsible for the Δ*gumB* virulence defect. At 24h post inoculation, the corneas infected with the Δ*gumB rcsB* double mutant had infiltrates similar to the WT, with 12 out of 12 infiltrates having a snow-flake like morphology (Figure 1A, C). At 48 h the corneal scores were indistinguishable between the Δ*gumB rcsC* and WT (Figure 2 and 3A), and this was reflected in restored IL-1β levels (p<0.05, ANOVA with Tukey’s posttest), (Figure 4B) the presence of hypopyons/hypopia (Figure 3C), and loss of corneal epithelia and numerous infiltrating neutrophils (Figure 5). CFU were similarly restored to the Δ*gumB* mutant by inactivation of the Rcs system (p<0.05 ANOVA with Tukey’s posttest, Figure 4C). Together, this data suggests that the Rcs system is a key mediator of *S. marcescens* corneal pathogenesis. Specifically, activation of the Rcs system reduces virulence and elimination of the Rcs system increases virulence to at least WT levels.

### *S. marcescens* gene expression during microbial keratitis is regulated by GumB

*S. marcescens* gene expression in the rabbit cornea was determined 24 hours post-infection using NanoString technology with a target set of 70 bacterial genes based on our previous studies, selected candidate virulence factors, and results of a preliminary RNA-Sequencing experiment performed using cultures grown in LB medium (to be described elsewhere). In two different experiments, eyes were infected with the Δ*gumB* mutant (n=4) and the WT (n=5). Of the 70 target genes, 39 were expressed above background levels by the WT while in the rabbit cornea (Table 3). Among the most highly expressed in the cornea included serralysin metalloproteases *prtS, slpB*, and *slpE*, TLR5-ligand flagellin, and importantly for this study genes from the Rcs system (*gumB* and *rcsB*) and the *shlA* cytolysin genes (Table 4).

**Table 3.**
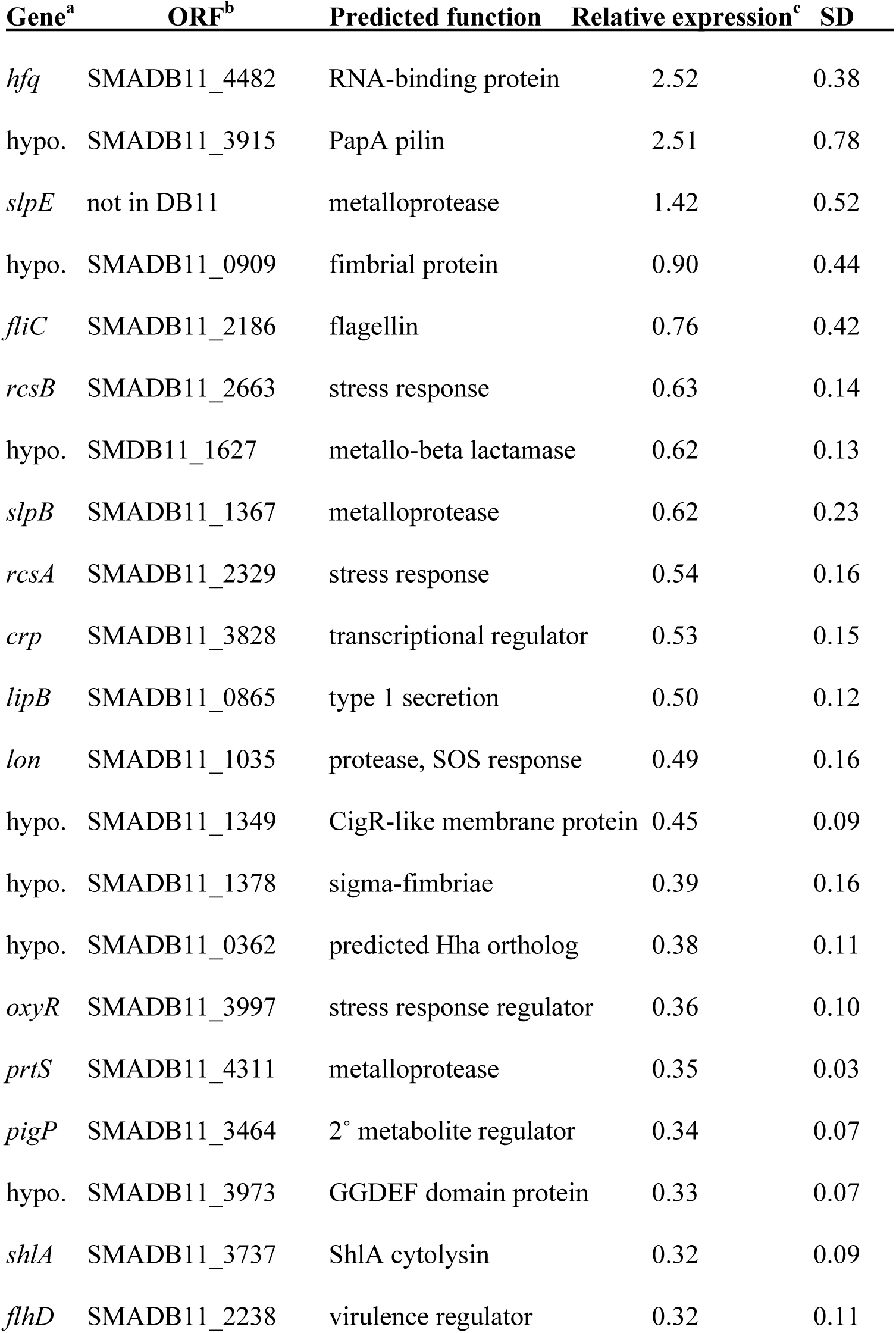
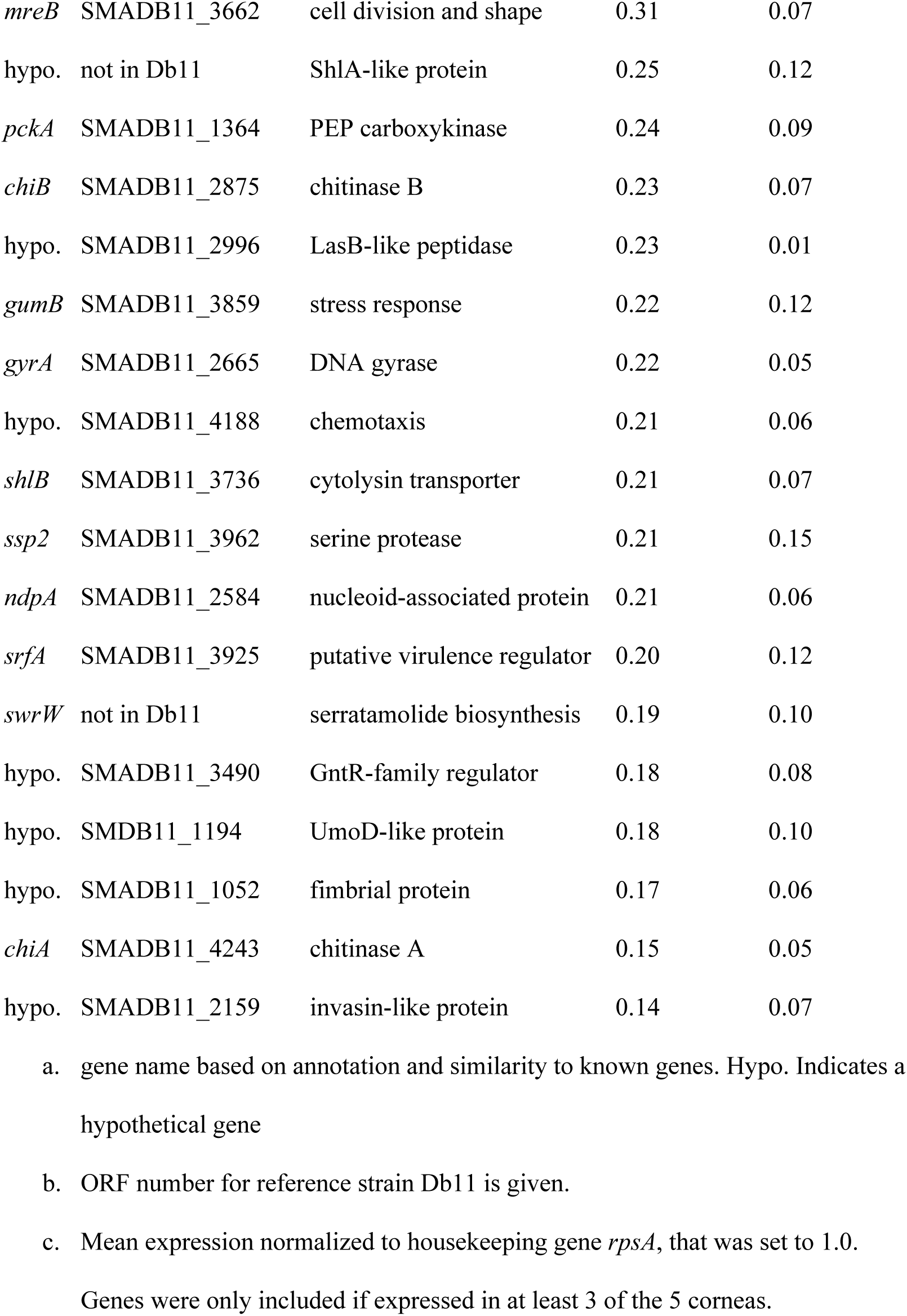
*S. marcescens* genes expressed bacterial during keratitis 24 h post-infection.

**Table 4.**
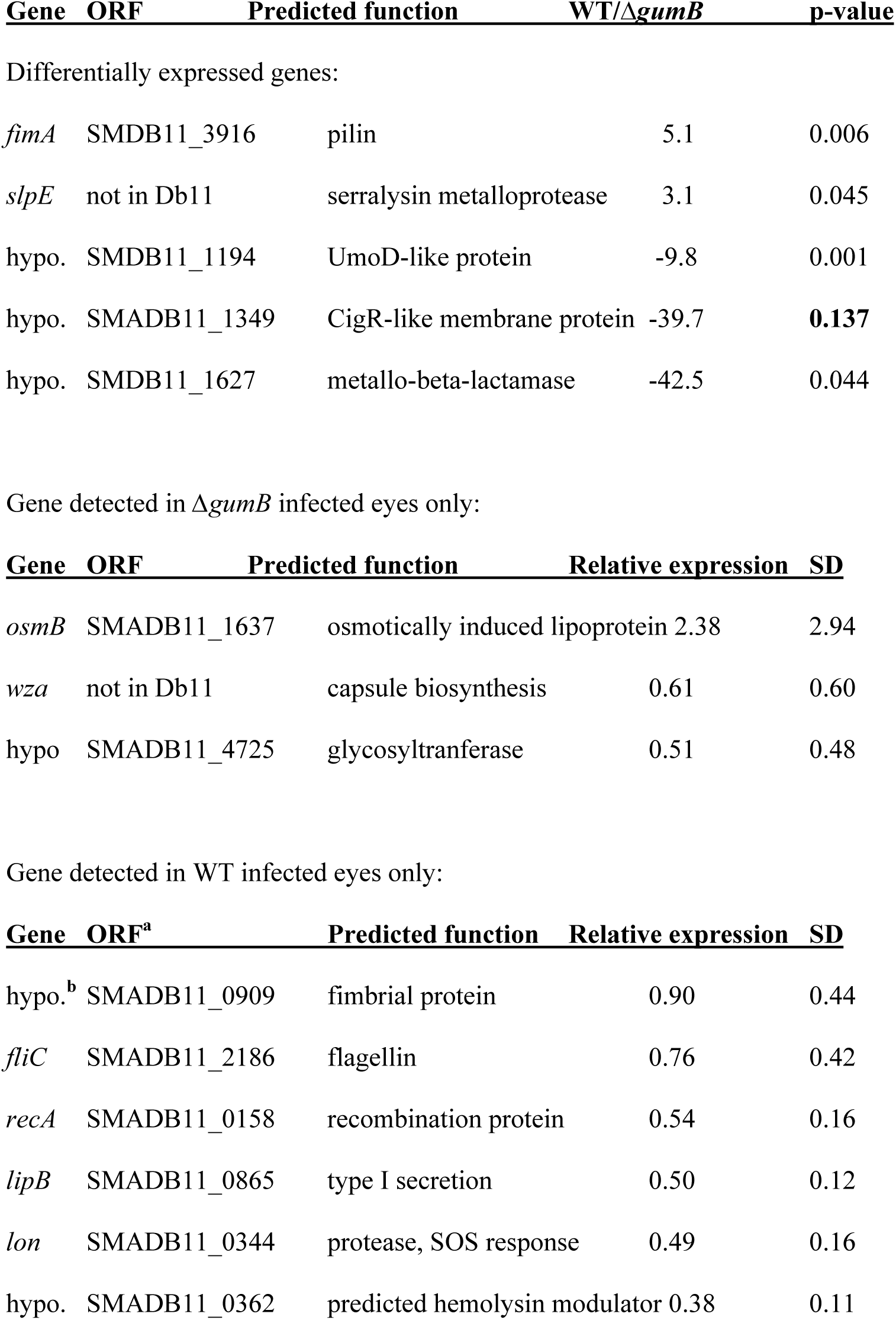
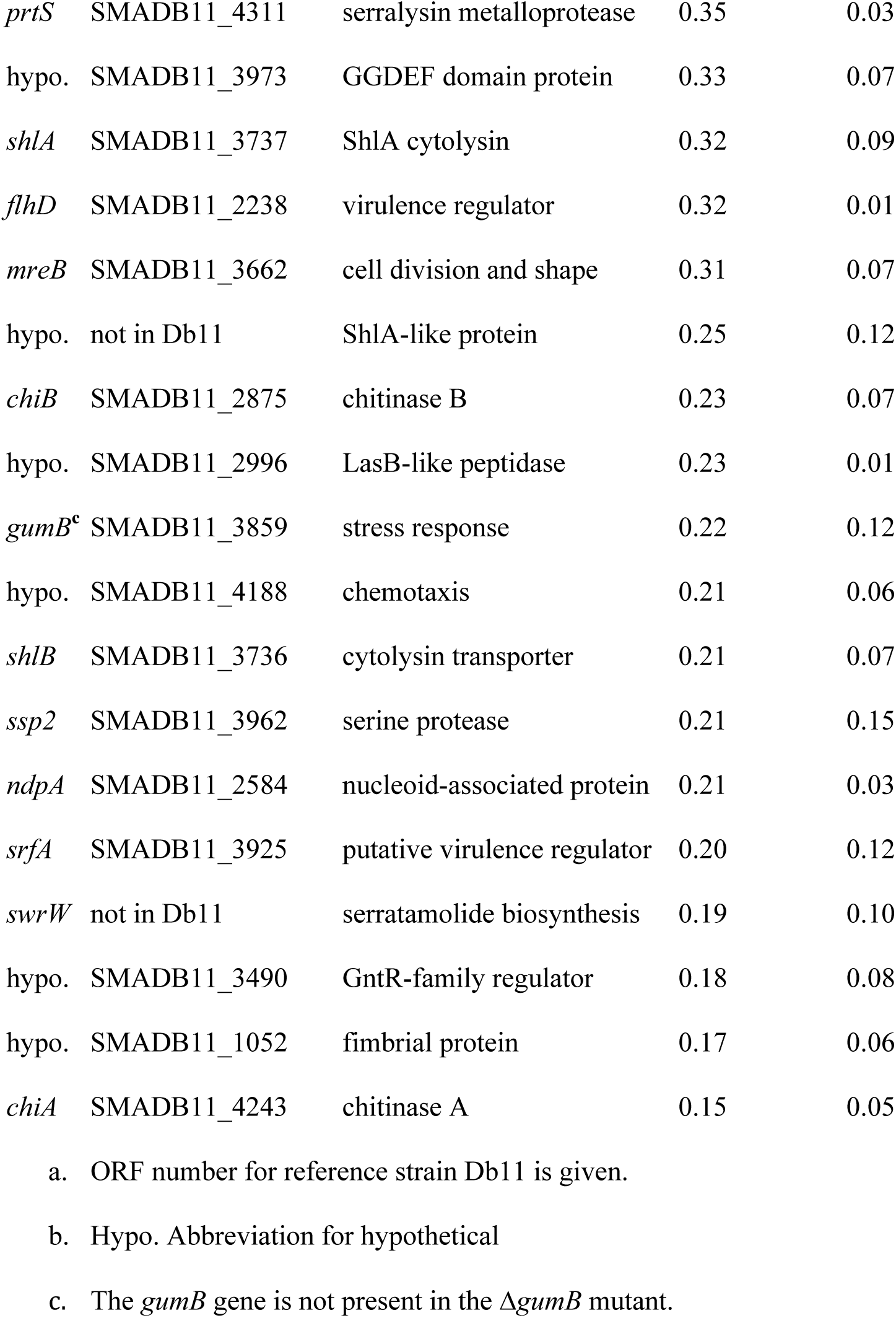
Differentially expressed bacterial genes during keratitis 24 h post-infection.

Twenty-four genes were measured from WT but not Δ*gumB* mutant infected eyes: these include motility related genes, a number of transcription factors, the serralysin protease and the a type I secretion system responsible for secretion of serralysin-family proteases, hemolysin associated genes including the *shlBA* operon, the *swrW* gene that codes for a nonribosomal peptide synthetase that generates the hemolytic biosurfactant serratamolide (also known as serrawettin W1) (Table 4). Conversely, three bacterial genes were detected from the Δ*gumB* infected eyes, but not WT infected eyes, including a capsule biosynthetic gene and a predicted osmotic stress induced lipoprotein (Table 4).

Ten other genes had a ± 2-fold change in expression between the WT and Δ*gumB* mutant. These include two genes with higher expression from the WT: serralysin-like metalloprotease SlpE and a type I pilus gene. The 8 genes expressed at higher levels from the *gumB* mutant include several stress related genes including *crp, oxyR*, and *rcsB* (Table 4). Importantly, these data verify that genes chosen in this study for analysis – those for the capsule, ShlA cytolysin, and Rcs system are expressed during corneal infections.

## DISCUSSION

Microbial keratitis is generally treatable using antibiotics, but still leads to vision loss due to rapid growth of the organism before treatment and continued damage to corneal tissue by inflammatory cells, even after the death of the microbes (23-27). In human patients, *S. marcescens* is a leading cause of contact-lens associated keratitis and can lead to vision loss and in rare cases, blindness (18, 19). *S. marcescens* has been shown to readily infect the corneas of rabbits (28-30). It is highly inflammatory, and its lipopolysaccharide and purified proteases are sufficient to induce keratitis (29, 31-34). In a mouse keratitis model, it was demonstrated that the cornea requires TLR4, TLR5, IL-1R and MyD88 to effectively clear *S. marcescens* infections (35), supporting that flagellin and LPS are important pathogen associated molecular patterns (PAMPS) of *S. marcescens* in the cornea. Nevertheless, the only *S. marcescens* gene tested for a role in keratitis using mutant bacterial strains thus far has been the transcription factor *eepR*, which was important for wild-type levels of bacterial proliferation within the rabbit cornea (36).

The purpose of this study was to establish whether the IgaA-family protein GumB from *S. marcescens* and the related Rcs stress response system play a role in microbial pathogenesis during bacterial keratitis. Data from this study demonstrated that a clinical keratitis isolate of *S. marcescens* with constitutively active Rcs system (Δ*gumB* mutant) was defective in causing corneal inflammation judged by corneal inflammatory signs, as the *gumB* mutant had a 62.5% reduction in corneal inflammatory scores. These correlate with reduced production of pro-inflammatory cytokines IL-1β and Tnfα and neutrophil infiltration into the cornea. Similarly, loss of *gumB* reduced the ability of the bacterium to proliferate in the cornea, with a 1.875-Log_10_ (98%) loss of CFU in *gumB* infected corneas relative to those infected by the WT. Complementation of the *gumB* mutation with wild-type *gumB* reversed the *gumB* mutant defects. Together, these data solidify the role for the Rcs system as having an important role in ocular bacterial host-pathogen interactions.

WT and the Δ*gumB* Δ*rcsC* strain, with an inactivated Rcs system, induced infiltrates with a snow-flake appearance at 24 h that were reminiscent of the presentation of infectious crystalline keratopathy, in which microbes are thought to form biofilms that spread along the lamelar planes of the corneal stroma that lead to a crystalline or snow-flake like pattern (37). This suggests that WT and Rcs defective strains penetrate into the cornea along the lamellar planes, whereas the Rcs-overexpressing Δ*gumB* mutant does not. The difference in spreading may be due to lack of swimming and swarming motility previously demonstrated for the Δ*gumB* mutant in *S. marcescens* and other members of the Enterobacteriaceae (9, 38, 39). The *in vivo* expression data presented here support the previous *in vitro* study that flagella and swarming motility-required genes, *fliC* and *swrW* are expressed by the WT but undetectable from the Δ*gumB* mutant. Additionally, expression data suggests highly reduced expression of bacterial proteases by the Δ*gumB* mutant. The *slpB* and *slpE* proteins are serralysins, a class of metalloproteases that have been implicated in the spread of *S. marcescens* in the cornea and necrosis and liquefaction of the cornea (28, 29, 40, 41).

Genetic experiments provided insight into the reduced pathogenesis of the Δ*gumB* mutant. Our results suggest that overexpression of the *S. marcescens* capsular polysaccharide contributes to the Δ*gumB* mutant phenotypes as revealed by intermediate corneal inflammatory phenotypes in the Δ*gumB* mutant in which the capsular polysaccharide genes *wecA* and *wza* were mutated (both being required for the extracellular capsule). However, the capsule was clearly not necessary for keratitis as the non-capsulated *gumB wecA* and *gumB wza* mutants both induced ulcers and proliferated in the cornea and caused more inflammation than the hyper-capsulated Δ*gumB* mutant. Notably, gene expression of the capsule was not measured in the WT bacteria during infection, further supporting the notion that it is not required for *S. marcescens* to cause keratitis in our model. In fact, the necessity of bacterial capsules during corneal infection has only been studied using isogenic strains with the Gram-positive bacterium *Streptococcus pneumoniae*. Marquart and colleagues demonstrated that *S. pneumoniae* has to be alive to cause ocular inflammation and that a capsule mutant was equally virulent to the capsulated wild type in a rabbit keratitis model (42, 43). In our study, the capsule may prevent the immune system from recognizing the bacteria surface PAMPS or by defending the bacteria against phagocytosis, leading to the reduced inflammation observed with the Δ*gumB* mutant.

A role was also evaluated for the ShlA cytolysin. ShlA, a key virulence factor for *S. marcescens* in several models, has not been tested in *in vivo* ocular models. Because expression of *shlA* and its transporter *shlB* is severely reduced in the *gumB* mutant, and *gumB* mutants generated a reduced keratitis relative to the wild type, testing a role for ShlA in the *gumB* keratitis defect was warranted. Here we observed that ectopic expression of *shlBA* in the *gumB* mutant partially restored corneal inflammatory signs of keratitis as well as proinflammatory cytokines. This occurred even though there was a clear selection for loss of the *shlBA* plasmid during the course of infection, relative to the empty vector. This suggests that the role of *shlBA* was underrepresented in this study, but the clear intermediate phenotypes of the *gumB* mutant with multicopy expression of *shlBA* supports the role for ShlA in ocular infections.

Whereas activation of the Rcs system through mutation of *gumB* led to reduced virulence, the inactivation of the Rcs system in the Δ*gumB rcsC* double mutant restored virulence to equal or higher than the WT. Similarly, in a bacteremia model *rcsB* mutants caused higher mortality of mice in a capsule-independent manner (6). Anderson and colleagues suggested that elevated *shlBA* expression in the *rcsB* mutant may be responsible for the increased virulence (6). This was concluded partially based on previous evidence describing that *rssAB* mutants, which have higher *shlBA* expression, caused higher mortality rates and more severe pathogenesis in a rat lung infection model (44). Together, these suggest that in *S. marcescens* the Rcs system may act to attenuate virulence, perhaps resulting in reduced immune activation and mortality of the host. Interestingly, species of the Enterobacteriaceae reported to have higher virulence in Rcs defective strains, all have ShlA orthologs (*E. tarda, P. mirabilis*, and *S. marcescens*), whereas those species reported to have reduced virulence conferred by *rcs* mutations typically do not have ShlA orthologs (*E. coli, C. rodentium* and *S. enterica*).

In a previous study the Δ*gumB* mutant was nearly completely avirulent using an invertebrate infection model with an approximately 4-Log_10_ reduction in CFU compared to the WT (11), whereas in the rabbit cornea they are reduced in proliferation, but just under 2-Log_10_ CFU. In both cases, however, the Δ*gumB* mutant was reduced in microbial pathogenesis, suggesting that the *G. mellonella* model was predictive of the mammalian model. The Δ*gumB* mutant has similar growth characteristics in minimal and rich media (9) and *G. mellonella* homogenates (11), suggesting that the observed defect is not due to reduced growth.

The constitutive expression of the Rcs system, through mutation of *gumB*, led to an overall reduction in bacterial CFU per cornea, but also highly reduced neutrophil infiltrates, correlating with reduced PAMP production and presentation. The bacteria developed into globular intrastromal infiltrates that are reminiscent of biofilms, which are typically more tolerant of host defenses and antibiotic therapy. The *in vivo* factors that activate the Rcs system are not fully understood, but include lysozyme and cationic peptides (3). Both of these are key features of the innate defenses of the ocular surface (45). It is likely that after infection is established, it would benefit *S. marcescens* to activate the Rcs system, which would shut off PAMP and toxin production and increase capsule production. However, during early stages, it is probable that some Rcs-inhibited factors contribute to establishing infection such as flagellin, ShlA, and serralysin family proteases. Thus, a well-coordinated balance in activity of the Rcs system is likely required for effective pathogenesis by *S. marcescens*. Consistent with this idea, the key Rcs system transcription factor, RcsB, is not only modulated in activity by its cognate histidine kinase, it also has numerous binding partners that fine tune its activity, such as RcsA (3). In summary, this study concludes the *S. marcescens* Rcs system is a major regulator of host-pathogen interactions during corneal infections.

## MATERIALS AND METHODS

### Bacterial strains and culture conditions

Bacterial strains are listed in Table 1 and were maintained at −80°C. Bacteria were streaked to single colonies and individual colonies were used to inoculate LB medium (46). Gentamicin was used at 10 µg/ml to select for plasmid maintenance.

The Δ*gumB rcsC* isolate (strain CMS4141), was isolated as a genetic suppressor of Δ*gumB* pigment phenotype (to be described elsewhere). The *rcsC* mutant gene was cloned in vector pMQ132 using primers and methods previously described (11). The mutation in *rcsC*, noted as *rcsC-SPV1*, was determined using Sanger sequencing of the *rcsC* gene (University of Pittsburgh Genomics Research Core).

### Microbial keratitis studies

The present study conformed to the ARVO Statement on the Use of Animals in Ophthalmic and Vision Research. The study was approved by the University of Pittsburgh Institutional Animal Care and Use Committee (IACUC Protocol 16098925B). For infection experiments, bacteria were grown overnight at 30°C with aeration, normalized by optical density (600 nm), and adjusted to ∼500 CFU in 25 µl, and injected into right eye of New Zealand White rabbits. Actual inocula colony counts were determined using the EddyJet 2 spiral plating system (Neutec Group Inc., Farmingdale, NY) on 5% trypticase soy agar with 5% sheep’s blood plates. The plates were incubated overnight at 30°C and the colonies were using the automated Flash and Grow colony counting system (Neutec Group). At 24 and 48 h post-injection, eyes were evaluated for ocular signs of inflammation using a slit-lamp according to a modified McDonald-Shadduck grading system (20). After sacrifice, corneas were harvested with a 10 mm trephine, homogenized with an MP Fast Prep-24 homogenizer using lysing matrix A tubes (MP Biomedicals), and bacteria were enumerated by dilution plating as described above. Homogenates were clarified by centrifugation (10 minutes at 16,000-21000 x g) and stored at −20°C until used for ELISA analysis for pro-inflammatory markers following manufacturers protocols and MPO analysis (47, 48). For IL-1β analysis, Sigma-Aldrich product RAB1108 was used, and for Tnfα measurement, R&D Systems DuoSet DY5670 was used.

RNA for the nanoString nCounter platform was harvested from corneas following the protocol of Xu, et al. (49) with a few exceptions. Corneas were removed with a trephine and immediately frozen in liquid nitrogen and stored at −80°C until RNA was harvested. Corneas were homogenized as described above. Flowing phenol chloroform extraction and ethanol precipitation, RNA was run through a ZymoResearch RNA concentrator column, adjusted to 800 ng/µl and sent to the University of Pittsburgh Genomics Research Core for nCounter analysis. Results were analyzed using nSolver software and results were uploaded to Gene Expression Omnibus (accession number GSE155486).

For histological analysis, corneas were fixed in 4% paraformaldehyde in phosphate buffered saline (pH 7.4) and embedded in paraffin. Central corneal sections were generated (5 µm) and stained with hematoxylin and eosin staining. Slides were imaged using the Olympus Provis AX-70 microscope with MagnaFire 2.1 software.

### Statistical analysis

Data was analyzed using GraphPad Prism Software. Non-parametric analysis was used to analyzed inflammatory scores by Mann-Whitney or Kruskal-Wallis analysis, and ANOVA with Tukey’s posttest or Student’s T-tests were used to analyze other experimental data.

## ACKNOWLEDGEMENTS

The authors thank Kimberly Brothers, Jake Callaghan, and Katherine Davoli for expert technical help, and Kira Lathrop, the Center for Biologic Imaging, Katherine Helfrich and Mark Ross for microscopy help. This study was supported by the Charles T. Campbell Laboratory of Ophthalmic Microbiology, NIH grant EY027331 (to R.M.Q.S.), and EY08098 (Core Grant for Vision Research), the Eye and Ear Foundation of Pittsburgh, and unrestricted funds from Research to Prevent Blindness.

